# Detection of stress-dependent m^5^C rRNA dynamics in *Escherichia coli* using m^5^C-Rol-LAMP

**DOI:** 10.1101/2025.06.13.659538

**Authors:** Leonardo Vásquez-Camus, Sebastian Riquelme-Barrios, Kirsten Jung

## Abstract

Numerous RNA modifications are known in prokaryotes, but their dynamics and function in regulation remain largely unexplored. In *Escherichia coli*, three methyltransferases catalyze the 5-methylcytosine (m^5^C) modification in ribosomal RNA. Here, we introduce m^5^C-rolling circle loop-mediated isothermal amplification (m^5^C-Rol-LAMP) as a novel qPCR-based method that offers high sensitivity and site-specific resolution to detect and quantify m⁵C in total RNA. When applying m^5^C-Rol-LAMP to *E. coli* under heat stress (45 °C), we observe a site-specific increase of m⁵C at position 1407 of 16S rRNA from 77% to 89%, while m^5^C levels at positions 967 (16S) and 1962 (23S) remain unchanged. In recovered cells (at 37°C), the m^5^C abundance partially returns to the no stress level. Under oxidative stress, the level of m⁵C1407 also increases, but remains high in recovered cells. These results demonstrate for the first time a reversible, stress-dependent and site-specific change in the rRNA modification level of a bacterium. m^5^C-Rol-LAMP is a powerful and easy-to-use tool for studying m^5^C in all RNA species, allowing the quantitative and site-specific detection of this modification.

## INTRODUCTION

The epitranscriptome, which includes over 180 RNA modifications identified to date (Boccaletto et al. 2022) that are deposited during or after transcription, influences gene transcription, RNA processing, nuclear export, cellular localization, and mRNA translation in eukaryotes (Shi et al. 2020; Delaunay et al. 2024). Among these modifications, methylation is one of the most common, found in all domains of life as well and in all RNA types, including mRNA, rRNA, and tRNA (Boccaletto et al. 2022). In particular, 5-methylcytosine (m^5^C) is one of the most prevalent modifications distributed along mRNAs, enriched in UTRs and conserved in tRNAs and rRNAs in eukaryotes (Chen et al. 2021). m^5^C is catalyzed by NSUN family enzymes and DNMT2 and plays a crucial role in RNA metabolism, including mRNA export, stability and translation (Bohnsack et al. 2019). Dysregulation of m^5^C modification or mutations in m^5^C methyltransferase genes are associated with various disorders, including diseases of the nervous system and cancer (Gao and Fang 2021). Although m^5^C has been extensively studied in eukaryotes, there is a large knowledge gap in prokaryotes.

In *Escherichia coli,* three methyltransferases responsible for m^5^C modification have been identified, each acting independently on specific sites within the rRNA: RsmB, RsmF, and RlmI. RsmB catalyzes the methylation of C967, and RsmF modifies C1407 in the 16S rRNA (Tscherne et al. 1999; Andersen and Douthwaite 2006). In contrast, RlmI is responsible for the methylation of C1962 in the 23S rRNA (Purta et al. 2008; Sunita et al. 2008). m^5^C is highly abundant in the mRNA of the hyperthermophile Archaeon *Thermococcus kodakarensis* (Fluke et al. 2024), but there is no evidence for the presence of m^5^C modification in *E. coli* mRNA (Edelheit et al. 2013; Riquelme-Barrios et al. 2025).

A systematic analysis of *E. coli* rRNA m^5^C methyltransferase knockout mutants revealed minor effects on ribosome assembly and protein production (Pletnev et al. 2020). RsmF methylates not only m^5^C1407 in the 16S rRNA of *E. coli*, but also tRNA in response to oxidative stress (Valesyan et al. 2024).

Detecting m^5^C in RNA poses several challenges. The most commonly used approach is bisulfite treatment, in which an unmethylated cytosine is selectively deaminated to uracil, while methylated cytosine remains unchanged (Gu et al. 2005; Schaefer et al. 2009). This treated RNA is then converted into a cDNA and sequenced, a method known as RNA bisulfite sequencing (RNA-BisSeq). Despite its utility, the method requires meticulous optimization and is limited by potential issues such as incomplete deamination (Motorin et al. 2010; Schaefer 2015). Liquid chromatography-tandem mass spectrometry (LC-MS/MS) offers another detection approach, but both methods have limitations in sensitivity and specificity (Guo et al. 2021). A modification of RNA-BisSeq allowed simultaneous detection of multiple RNA modifications, including m^5^C, at single base resolution. However, this method detected significantly fewer m^5^C sites in both coding and non-coding RNAs compared to previous studies (Khoddami et al. 2019). The limitations of RNA-BisSeq combined with the requirement for sophisticated equipment and the labor-intensive sequencing underscore the urgent need for sensitive, accurate and cost-effective methods to comprehensively study m^5^C in various RNA species.

Using RNA direct sequencing (DRS), our group recently studied the epitranscriptomic response of *E. coli* to heat stress (45°C) (Riquelme-Barrios et al. 2025). The mass spectrometry (MS) analysis revealed a 5% increase in m^5^C abundance under heat stress, however; the specific site contributing to this increase remains unidentified (Riquelme-Barrios et al. 2025). To investigate the dynamics of m^5^C modification during stress response, we developed m^5^C-Rol-LAMP, a novel isothermal qPCR-based technique for the site-specific detection and quantification of m^5^C. Applying m^5^C-Rol-LAMP, we uncovered that 16S rRNA m^5^C1407 modification in *E. coli* increases under heat stress and returns to almost the no stress level during recovery at 37°C. Extending the analysis to oxidative stress revealed an increase at the same m^5^C site which remained high in recovered cells.

## RESULTS

### m^5^C-Rol-LAMP is able to detect m^5^C at known single-nucleotide positions in the rRNA of *E. coli*

Based on the rolling circle extension-actuated loop-mediated isothermal amplification (RCA-LAMP) for microRNAs (Tian et al. 2019) and rolling circle extension-assisted loop-mediated isothermal amplification for m^6^A (m^6^A-Rol-LAMP) (Li et al. 2023), we have developed a new approach for the detection and quantification of m^5^C (**Figure 1**). The method first uses bisulfite treatment of total RNA to distinguish m^5^C from unmodified cytosine. In this process, unmodified cytosines are converted to uracil, while m^5^C is retained as cytosine (**Figure 1A**). Then, a padlock probe that contains a backbone for the rolling circle amplification followed by a loop-mediated isothermal amplification (RCA-LAMP) (**Supplementary Table 1**), is hybridized to the target RNA molecule (**Figure 1B**). The 5’ and 3’ arms of the probe are complementary to the flanking regions of an m^5^C site of interest, leaving a single nucleotide gap (**Figure 1B**). To close the gap, a polymerization/ligation step is done by using DNA polymerase BST 2.0 and SplintR Ligase that efficiently catalyzes the ligation of adjacent single-stranded DNA splinted by a complementary RNA strand (Jin et al. 2016). In this step, two independent reactions are performed: one with dATP and one with dGTP (**Figure 1C**). The gap site of the RNA-DNA padlock probe complex is filled with dATP or dGTP, depending on whether the analyzed cytosine position was previously modified or not (dGTP for modified cytosines and dATP for unmodified cytosines). Once the gap is closed, amplification occurs in a single-step process (**Figure 1D**) by combining the circularized padlock probes with a forward internal primer (FIP), backward internal primer (BIP) and stem-loop primer (SLP) primer (**Supplementary Table 2**). Briefly, FIP initiates a rolling circle amplification by binding to the circular padlock probe and produces a long linear ssDNA. This ssDNA forms a double stem-loop DNA via SLP binding, which triggers loop-mediated amplification. The amplification products can be detected by real-time PCR (Li et al. 2023).

**Figure 1:**
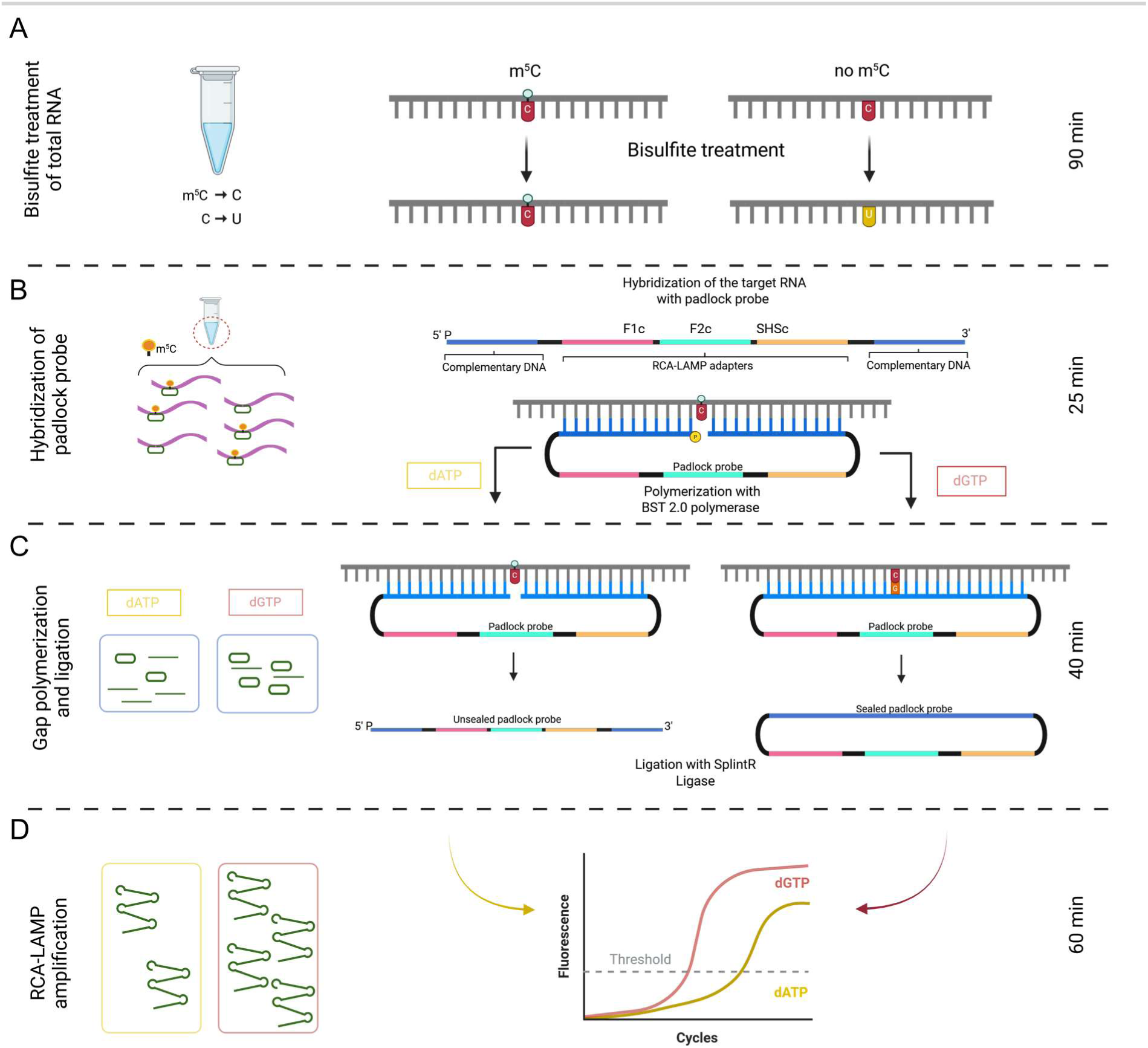
Schematic representation of m^5^C-Rol-LAMP. Workflow of the m^5^C-Rol-LAMP technique including time management. A) Bisulfite treatment of total RNA. B) Hybridization of the padlock probe. C) Gap polymerization and ligation. D) RCA-LAMP amplification, measured by fluorescence. Error bars represent standard deviations of the mean of three replicates. * P < 0.05; ** P < 0.01; *** P < 0.001; **** P < 0.0001; by unpaired two-tailed t-test. position. B) Ct values were plotted against the logarithm of the input concentration. Reactions were performed with either dATP, dGTP, or dATP+dGTP as ligation substrates. The addition of dATP+dGTP results in maximum amplification and serves as control. The R² values indicate high linearity of the assays (dATP R² = 0.9938; dGTP R² = 0.9940; dATP+dGTP R² = 0.9940). Error bars represent standard deviations of the mean of three replicates. * P < 0.05; ** P < 0.01; *** P < 0.001; **** P < 0.0001; n.s. not significant by unpaired two-tailed t-test.

For proof of concept, we used this technique to detect the known m^5^C positions in the rRNA of wild-type *E. coli* compared to an m^5^C depleted mutant. For this purpose, we generated a triple mutant (Δ*rsmB*Δ*rsmF*Δ*rlmI*) lacking the methyltransferases that catalyze m^5^C methylation at positions 1962 of 23S rRNA (**Figure 2A**), 967 of 16S rRNA (**Figure 2B**) and 1407 of 16S rRNA **(Figure 2C)**. In the wild-type strain, where m^5^C is present (Riquelme-Barrios et al. 2025; Petrov et al. 2022), the threshold cycle (Ct) values were lower with dGTP than with dATP, indicating that more amplification products were generated with dGTP because the modified cytosine remained as cytosine after bisulfite treatment. In the mutant, Ct values were lower with dATP than with dGTP, indicating that more amplification products were generated with dATP because all cytosines were converted to uracil after bisulfite treatment. By comparative analysis of wild type and mutant by m^5^C-Rol-LAMP assays, with dATP or dGTP, we detected m^5^C methylation at all known positions in *E. coli* rRNA (**Figure 2A-C**).

**Figure 2:**
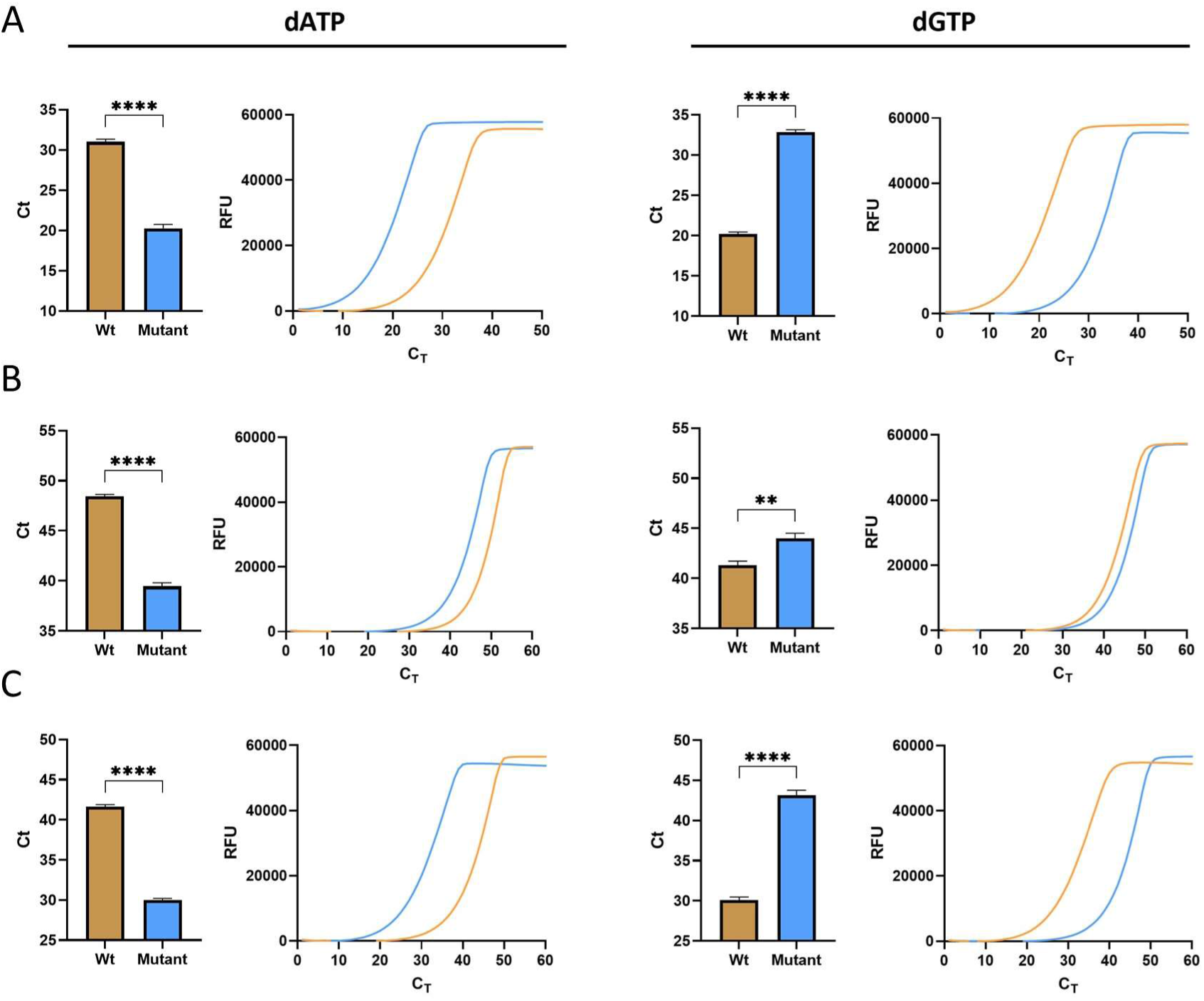
Detection of m5C rRNA modification at the known sites in total RNA of E. coli MG1655 and E. coli ΔrsmBΔrsmFΔrlmI mutant with m5C-Rol-LAMP. Bar graph and amplification curves for 23S rRNA m5C1962 (A), 16S rRNA m5C967 (B) and 16S rRNA m5C1407 (C). RFU – relative fluorescence units. Error bars represent standard deviations of the mean of three replicates. * P < 0.05; ** P < 0.01; *** P < 0.001; **** P < 0.0001; by unpaired two-tailed t-test.

To evaluate the sensitivity of m⁵C-Rol-LAMP, we tested serial dilutions of a 1:1 mixture of methylated and unmethylated m^5^C synthetic RNA oligonucleotides. These synthetic RNA oligos corresponded to the native sequence surrounding the 23S rRNA m⁵C1962 site (details in Materials and Methods). The method reliably quantified methylation up to 0.1 fmol RNA input (**Figure 3A**). Ct values were similar for the dATP and dGTP reactions as it was as expected for a 50% methylated input. The assay with a mixture of dATP+dGTP allowed ligation at both modified and unmodified sites and could be used as control value for maximum padlock probe amplification (**Figure 3A**). Ct values showed strong linearity over the dilution range (R² > 0.99) (**Figure 3B**).

**Figure 3:**
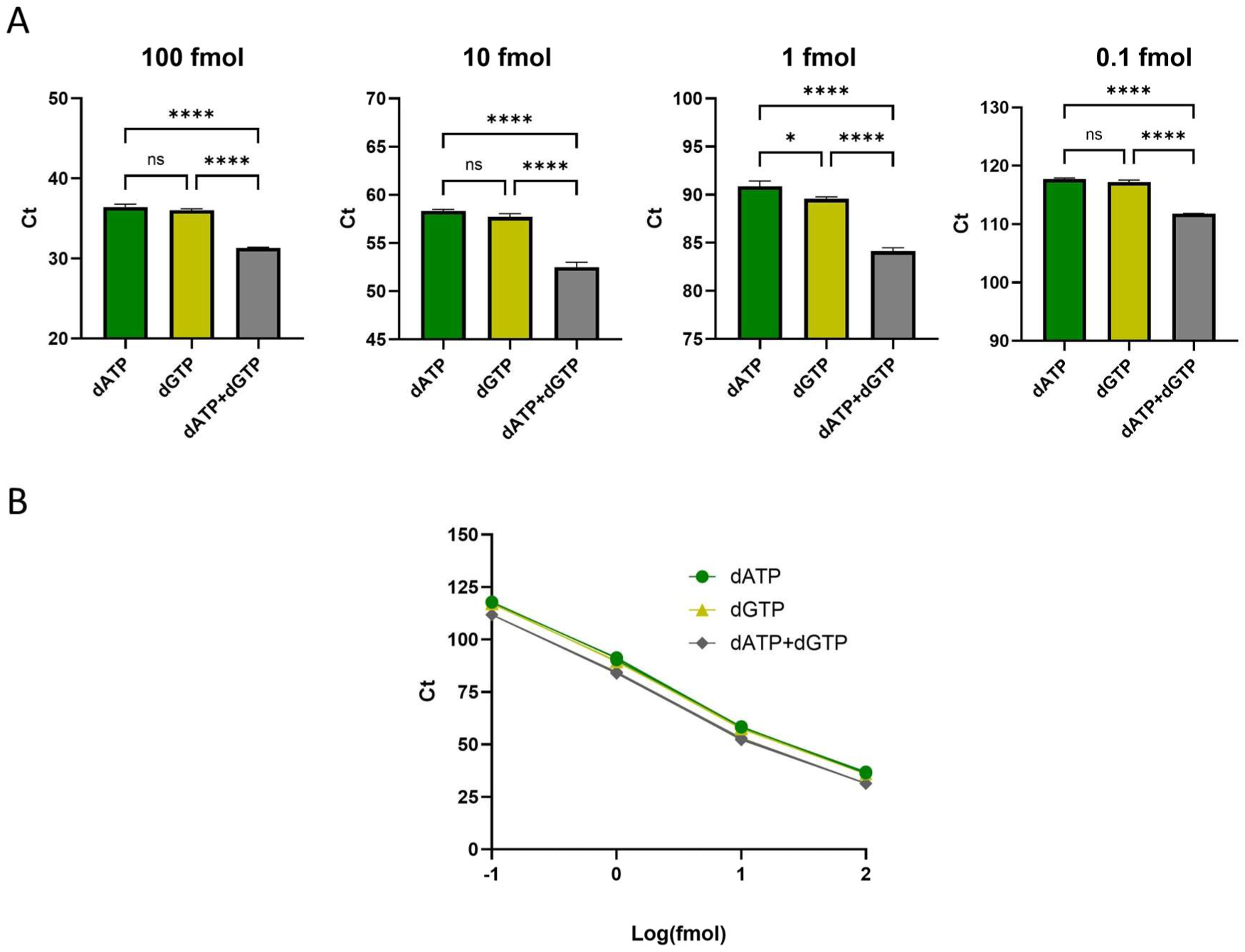
Sensitivity of the m⁵C-Rol-LAMP assay using synthetic RNA oligos mimicking the 23S rRNA m⁵C1962 site. A) Bar graphs of threshold cycles using serial dilutions (100 fmol to 0.1 fmol) of a 1:1 mix of synthetic RNA oligos containing methylated or unmodified cytosine at the 23S rRNA m⁵C1962 position. B) Ct values were plotted against the logarithm of the input concentration. Reactions were performed with either dATP, dGTP, or dATP+dGTP as ligation substrates. The addition of dATP+dGTP results in maximum amplification and serves as control. The R² values indicate high linearity of the assays (dATP R² = 0.9938; dGTP R² = 0.9940; dATP+dGTP R² = 0.9940). Error bars represent standard deviations of the mean of three replicates. * P < 0.05; ** P < 0.01; *** P < 0.001; **** P < 0.0001; n.s. not significant by unpaired two-tailed t-test.

### The methylation level of 16S rRNA m^5^C1407 changes during heat stress in *E. coli*

Mass spectrometry (MS) has previously identified a 5% increase in m^5^C abundance in *E. coli* rRNA under heat stress (45°C) (Riquelme-Barrios et al. 2025). However, this approach showed an overall increase and could not determine which of the three known rRNA m^5^C sites was responsible for this change. Here, we used three site-specific padlock-probes and analyzed the degree of methylation at each position in the total RNA of unstressed cells, stressed cells and recovered cells with m^5^C-Rol-LAMP (**Figure 4A**). *E. coli* MG1655 was grown in LB medium at 37°C to an OD600 of 0.5, and the “no stress” sample was collected. Hot LB medium was then added to immediately raise the temperature to 45°C, followed by incubation at 45°C for 30 min, and the “heat stress” sample was collected (Riquelme-Barrios et al. 2025). Cold LB medium was then added to immediately decrease the temperature to 37°C, followed by incubation at 37°C for 30 min and the “recovery” sample was collected (**Figure 4A**). Every condition was tested performing two m^5^C-Rol-LAMP reactions: one with dATP (**Figure 4B**) and one with dATP+dGTP (**Figure 4C**). In the presence of dATP, gap polymerization and ligation occur at uracil (unmethylated cytosine converted to uracil by bisulfite treatment) and thus represents the unmethylated fraction. In the presence of dATP+dGTP, gap polymerization and ligation occur on both, the methylated and unmethylated cytosines, and represents maximal padlock probe amplification as a transcript abundance control.

**Figure 4:**
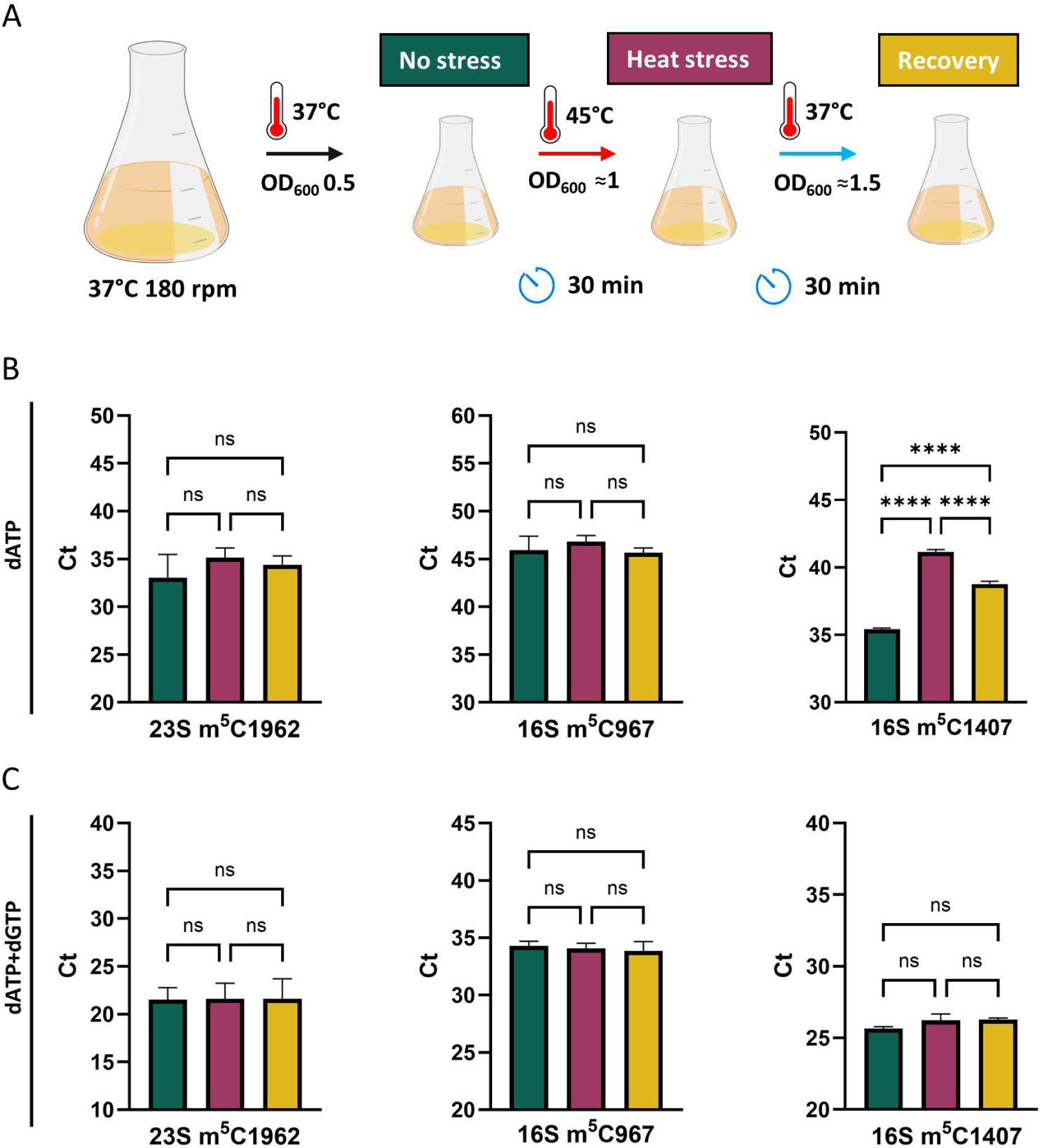
m^5^C rRNA modification in *E. coli* under heat stress. A) Workflow of the experiment. *E. coli* MG1655 was cultivated at 37°C until an OD600 of 0.5 was reached, and the first sample was collected (no stress, green). Heat stress was induced by adding hot LB medium to immediately increase the temperature to 45°C. After 30 min of incubation the second sample (heat stress, red) was collected. Cold LB medium was added to immediately lower the temperature to 37°C and after 30 min the third sample (recovery, yellow) was collected. B-C) Total RNA was analyzed by m^5^C-Rol-LAMP using the indicated site-specific padlock probes with dATP (B) and dATP+dGTP (C) in the ligation step. Shown are the mean Ct values of three biological replicates. Error bars indicate the standard deviation of the mean. * P < 0.05; ** P < 0.01; *** P < 0.001; **** P < 0.0001; n.s. not significant by One-way ANOVA with Tukey’s post hoc test.

By comparing the Ct values of these two reactions, we were able to detect differences between the three methylation sites. We found a significant increase in the Ct value for 16S rRNA m^5^C1407 in the dATP reaction during heat stress compared to the no stress condition, indicating that uracil was less abundant, implying an increase in m^5^C abundance at this specific site (**Figure 4B**). In the recovered cells, the Ct value for 16S rRNA m^5^C1407 was significantly reduced, but did not reach the no stress level, indicating a reversible response. It is important to note that Ct values of the dATP+dGTP reactions did not change under all conditions confirming equal rRNA amounts in the three samples **(Figure 4C)**. In contrast, Ct values determined in the presence of dATP or dATP+dGTP for the other two m^5^C modification sites remained unchanged under all conditions (**Figure 4B, C**). These results demonstrate for the first-time locus-specific, reversible stress-dependent dynamics of the m^5^C level of 16S rRNA in *E. coli*.

### The methylation level of 16S rRNA m^5^C1407 increases in response to oxidative stress in *E. coli*

Previously, an increase of the m^5^C level in Tyr-QUA II tRNA was reported under oxidative stress imposed by H2O2 in *E. coli* (Valesyan et al. 2024). We used m^5^C-Rol-LAMP to test whether the m^5^C methylation level also changes at the rRNAs (**Figure 5**). *E. coli* MG1655 was grown in LB medium at 37°C to an OD600 of 0.5, and the “no stress” sample was collected. 2 mM of hydrogen peroxide was then added. After 15 min, the “ox stress” sample, and after 60 min the “recovery” sample was collected (**Figure 5A**). Similar to heat stress, the Ct value of 16S rRNA m^5^C1407 of the dATP reaction increased during oxidative stress, and it remained high even when cells continued to grow (recovery phase) (**Figure 5B**). The Ct values of the dATP+dGTP reaction remained constant under all conditions, which confirmed that the determined effect was not related to differences in RNA quantities (**Figure 5C**). Notably, the other two known m^5^C sites in *E. coli* rRNA (16S rRNA m^5^C967 and 23S rRNA m^5^C1962) remained unchanged in the dATP and dATP+dGTP reactions (**Figure 5B, C**). These results reveal a locus-specific increase of the m^5^C level at position m^5^C1407 of 16S rRNA under oxidative stress.

**Figure 5:**
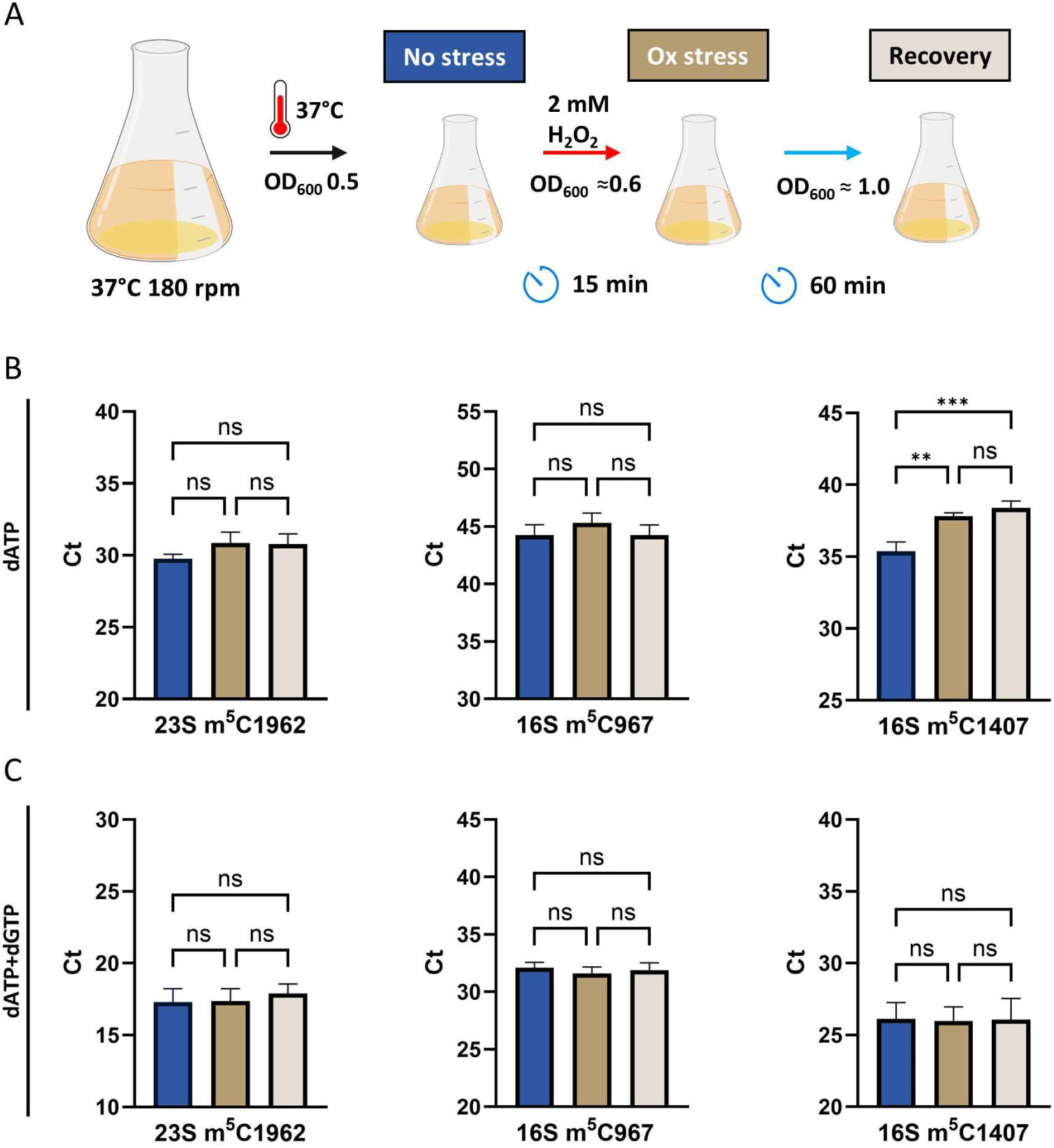
m^5^C rRNA modification in *E. coli* under oxidative stress. A) Workflow of the experiment. *E. coli* MG1655 was cultivated at 37°C until an OD600 of 0.5 was reached, and the first sample was collected (no stress, blue). Oxidative stress was induced by adding 2 mM H2O2 and cultivation was continued. Samples were collected after 15 min (oxidative stress, brown) and 60 min (recovery, light brown). B-C) Total RNA was analyzed by m^5^C-Rol-LAMP using the indicated site-specific padlock probes with dATP (B) and dATP+dGTP (C) in the ligation step. Shown are the mean Ct values of three biological replicates. Error bars indicate the standard deviation of the mean. * P < 0.05; ** P < 0.01; *** P < 0.001; **** P < 0.0001; n.s. non-significant by One-way ANOVA with Tukey’s post hoc test.

### Quantitative measurement of 16S rRNA m^5^C1407 levels confirms reversible dynamics under heat stress and an increase under oxidative stress

While the shift in Ct values for 16S m⁵C1407 during heat stress and oxidative stress provided a qualitative indication of increased modification, we sought to quantify this change. To do so, we utilized the quantitative capabilities of m⁵C-Rol-LAMP by calibrating amplification output against known amounts of synthetic RNA oligos (**Supplementary Table 3**), each ≈ 40 nucleotides in length and containing either a methylated (m⁵C) (**Figure 6A**) or an unmethylated cytosine (data not shown) centrally positioned to mimic the native sequence context around position 1407 of the *E. coli* 16S rRNA. Ligation reactions were performed with dATP+dGTP to obtain the maximum number of effectively closed padlock probes. The Ct values were plotted against the logarithmic values of the amount of input RNAs resulting in a linear correlation. We used this calibration curve to convert Ct values from our RNA samples into amounts of effectively closed padlock probes. By comparing reactions with dATP (which detects unmethylated cytosines converted to uracil) to dATP+dGTP (which detects unmethylated and methylated cytosines), we were able to quantify the proportion of methylated cytosines in our samples. In non-stressed cells, the methylation level at 16S rRNA m⁵C1407 was 77.16% ± 0.65. Under heat stress, the methylation increased to 89.51% ± 0.36 indicating an increase of approximately 12% (**Figure 6B**). After recovery, the methylation level decreased to 83.40% ± 0.80 but remained at a higher level compared to the non-stressed cells. These results demonstrate a dynamic regulation of 16S rRNA m⁵C1407 methylation in *E. coli* in response to heat stress, which is characterized by a rapid increase followed by a partial reset upon return to non-stress conditions.

**Figure 6.**
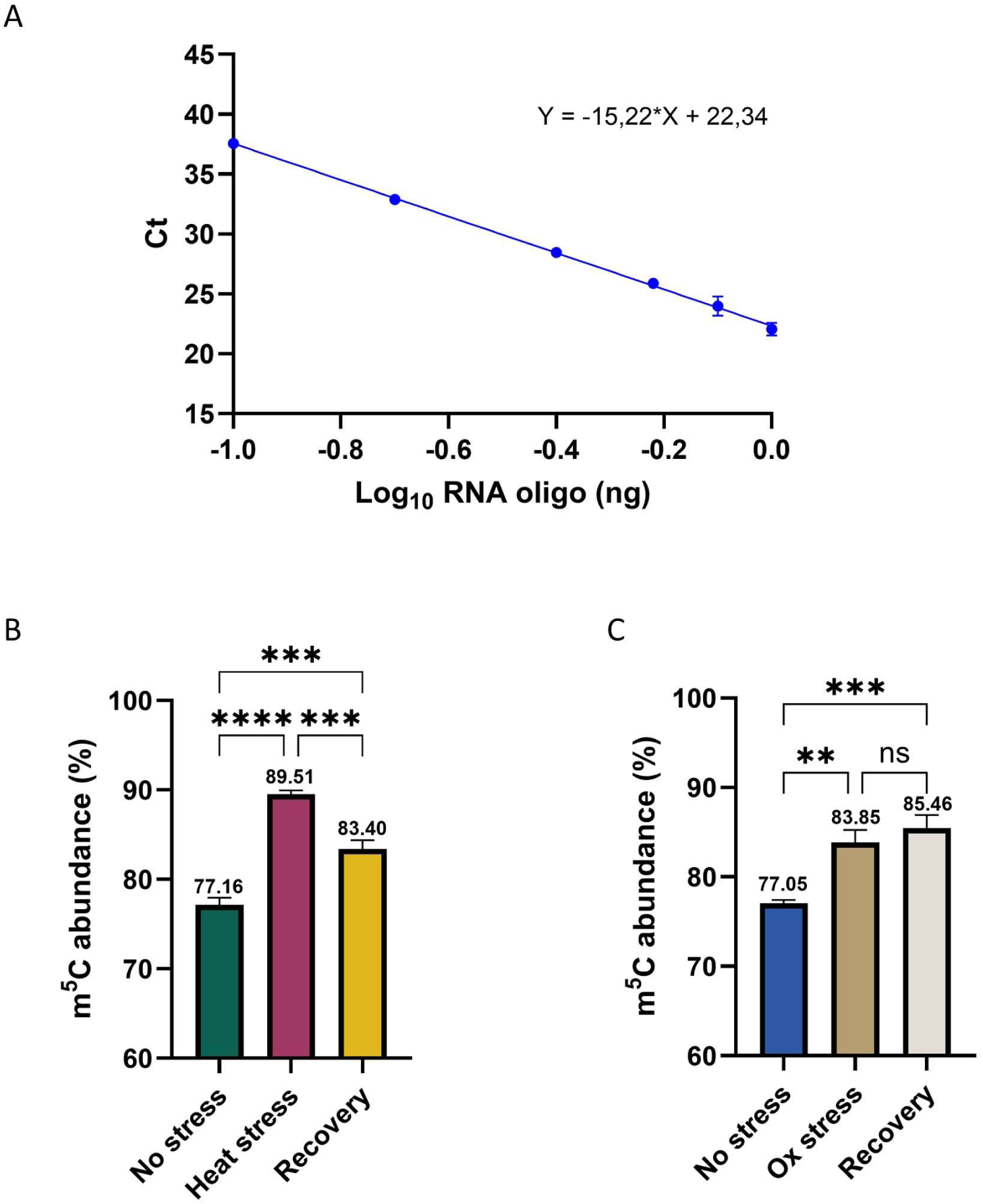
Quantification of the degree of methylation of 16S rRNA m⁵C1407 using m⁵C-Rol-LAMP. A) A calibration curve was generated using m⁵C-Rol-LAMP by amplifying increasing amounts of a synthetic 41-nucleotide RNA oligo representing the sequence around position 1407 of *E. coli* 16S rRNA with a methylated cytosine. Reactions were performed using dATP+dGTP to achieve maximum amplification of all correctly hybridized padlock probes. Ct values were plotted against the logarithmic values of the RNA inputs [RNA oligo (ng)]. Each data point represents the average of three independent replicates and error bars indicate the standard deviation of the mean. B) Quantification of m⁵C1407 methylation levels in *E. coli* under no stress, heat stress, and recovery conditions. C) Quantification of m⁵C1407 methylation levels under no stress, oxidative stress and recovery conditions. Methylation percentages were interpolated using the Ct values from dATP reactions (unmethylated fraction) relative to those from dATP+dGTP reactions (unmethylated and methylated cytosine), using the calibration curve in (A). Shown are the mean values of three biological replicates. Error bars indicate the standard deviation of the mean. * P < 0.05; ** P < 0.01; *** P < 0.001; **** P < 0.0001; n.s. non-significant by One-way ANOVA with Tukey’s post hoc test.

In the case of oxidative stress (**Figure 6C**), the methylation level was determined with a 77.05% ± 0.31 under non-stress conditions. Under oxidative stress, this value increases to 83.85% ± 1.15 after 15 min and remained elevated (86.46% ± 1.19) after 60 min. This sustained increase suggests that 16S rRNA m⁵C1407 methylation not only responds to stress but may also play a role in the recovery phase.

## DISCUSSION

Here we present m^5^C-Rol-LAMP as a new qPCR-based method for the detection and quantification of locus-specific changes in m^5^C methylation in RNA. Other techniques such as RNA-BisSeq, m^5^C-RIP (Gu and Liang 2019), miCLIP (George et al. 2017), MS-LS focus mainly on the identification/detection of modification sites and overall changes on a given sample, but they lack the sensitivity to quantify changes at the nucleotide specific level in a time and equipment-efficient manner (**Figure 1**). The ability of m^5^C-Rol-LAMP to quantitatively detect changes in m^5^C abundance underscores its usefulness for epitranscriptome research.

We used this technique to determine the site-specific m^5^C modification level of rRNA in *E. coli* under stress. We were able to demonstrate that one of the three reported modified sites, 16S rRNA m^5^C1407, shows a significant increase in abundance upon heat stress, which diminishes in recovered cells. This observation suggests that 16S rRNA m^5^C1407 plays a specific role in *E. coli* in the adaptation to higher temperatures. Under oxidative stress, we observed a similar increase in 16S rRNA m^5^C1407 abundance, which remained high in recovered cells. These results reveal that m⁵C1407 is crucial for the fine-tuned stress response of *E. coli*. m⁵C1407 is located in the decoding center of 16S rRNA, a structurally and functionally critical region of the ribosome. It should be noted that other nearby modifications, such as m⁴Cm1402 and m⁶,⁶A1518–1519, also show a heat stress-induced increase in abundance (Riquelme-Barrios et al. 2025). We speculate that this collective increase in rRNA modifications around the decoding center may serve as a protective and/or stabilizing response of the ribosome under heat stress and oxidative stress to preserve decoding fidelity and translation efficiency under stress, especially when the modifications of tRNAs have also changed (Riquelme-Barrios et al. 2025). Previous results indicated that m^5^C methylation is crucial for the adaptation of the thermophilic *T. kodakarensis* to high temperature (Fluke et al. 2024).

The reversible nature of methylation at m⁵C1407 raises the possibility of an active demethylation mechanism, an "eraser" enzyme, in bacteria. While demethylases are well described in eukaryotes (Shi et al. 2019), no such enzyme has yet been identified in prokaryotes. It is important to note that the level of m^5^C methylation at all three positions is relatively high in unstressed cells. Therefore, only a fraction (about 23%) of ribosomes can be methylated at position m⁵C1407 16S rRNA under heat or oxidative stress, so methylation dynamics could also be due to ribosomal heterogeneity and/or single cell heterogeneity to optimally adjust the translation rate of cells under stress (Pavlou et al. 2025).

In summary, m⁵C-Rol-LAMP enables the site-specific and quantitative analysis of m^5^C RNA modification. While this study focuses on m^5^C in *E. coli*, the broader applicability of m^5^C-Rol-LAMP to other organisms and RNA remains an exciting avenue for future research, and reinforce the idea of RNA modification as key player in stress response regulation. Notably, m^5^C has gained significant attention as a key regulator in tumorigenesis and cancer immunotherapy (Song et al. 2022). This method is inexpensive, and the ease of analysis using a qPCR-based technique makes m^5^C-Rol-LAMP an ideal approach for studying the changes and dynamics of m^5^C RNA modifications in both basic research and clinical settings.

## MATERIALS AND METHODS

### Bacterial strains and growth conditions

Experiments were conducted using the *Escherichia coli* strain MG1655 and the *E. coli* MG1655 *ΔrsmBΔrsmFΔrlmI* mutant. Bacteria were grown in lysogeny broth (LB) at 37°C with continuous shaking at 180 rpm to an optical density at 600 nm (OD600) of 0.5. The *E. coli* mutant was generated with in-frame deletion using an approach similar to a previously described (Lassak et al. 2010). Briefly, chromosomal deletions were generated using a two-step recombination approach based on a suicide vector system. Approximately 500 bp sequences flanking the target gene were amplified by PCR and assembled into the non-replicative plasmid pNPTS138-R6KT using Gibson assembly. The construct was first introduced into *E. coli* DH5α λpir for propagation, then transferred into the *E. coli* WM3064 donor strain, which requires diaminopimelic acid (DAP) for growth. Conjugative transfer into *E. coli* MG1655 was performed on LB agar supplemented with 300 μM DAP to facilitate genomic integration. Kanamycin-resistant colonies were selected for chromosomal integration of the plasmid, followed by counter-selection on sucrose-containing medium (10% w/v) to isolate double-crossover events. Colonies resistant to sucrose and sensitive to kanamycin were screened by colony PCR and verified by sequencing to confirm gene deletion. All primers used are listed in **Supplementary Table S4**.

### Oligo design, padlock probes, and primers

The sequences of padlock probes, RCA-LAMP specific primer and synthetic RNA oligos are listed in **Supplementary Tables 1,2 and 3**. RNA oligos were designed to target three known methylated cytosine positions in *E. coli* rRNA: 23S m⁵C1962, 16S m⁵C967, and 16S m⁵C1407. For each site, RNA oligonucleotides of ≈ 40 nucleotides in length were synthesized to mimic the native sequence context surrounding the modified cytosine, which was placed centrally to ensure accurate representation of structural and sequence features influencing ligation efficiency. For quantification, matched pairs of synthetic oligos containing either a methylated cytosine (m⁵C) or an unmodified cytosine at the target position were used. Modified and non-modified oligos used in this study were synthesized by Integrated DNA Technologies (IDT, Leuven, Belgium).

Padlock probes were designed to hybridize immediately upstream and downstream of the methylation site, flanking the central cytosine. Probe arms were selected to ensure optimal melting temperature (Tm), minimize secondary structures, and avoid off-target binding. Ligation was designed to occur directly at the modification site, enabling discrimination between methylated and unmethylated templates in the presence of selective nucleotide combinations during the ligation step (dATP, dGTP or dATP+dGTP). Padlock probes and primers were synthesized by Merck (Merck KGaA, Darmstadt, Germany).

### Heat stress

*E. coli* MG1655 was grown in LB medium at 37°C to an OD600 of 0.5. To induce heat stress, 7.11 mL of pre-warmed LB (90°C) was added to 40 mL of the bacterial culture (OD600 = 0.5) resulting in a final temperature of 45°C. The culture was then incubated at 45°C for 30 min under continuous shaking at 180 rpm. Then, 7.27 mL of chilled LB (4°C) was added to 30 mL of the 45°C culture to lower the temperature to 37°C. The culture was then incubated at 37°C for an additional 30 min with shaking at 180 rpm. Cell samples (10 mL) were collected at three time points: immediately before heat stress (no stress), after 30 min of incubation at 45°C (heat stress), and after 30 min recovery at 37°C (recovery).

### Oxidative stress

*E. coli* MG1655 was grown in LB medium (50 ml) at 37°C with shaking at 180 rpm to an OD600 of 0.5.

Hydrogen peroxide was added to a final concentration of 2 mM to induce oxidative stress and cultivation was continued at 37°C for 60 min under continuous shaking at 180 rpm. Cell samples (10 mL) were collected at three time points: immediately before hydrogen peroxide addition (no stress), 15 min after peroxide treatment (oxidative stress), and 60 minutes after peroxide treatment (recovery).

### Total RNA extraction

RNA extraction was performed using the PCI protocol (Sambrook and Russell 2006) with modifications (Petrov et al. 2022). Briefly, bacterial cultures were combined with phenol and ethanol to achieve final concentrations of 1% (v/v) and 20% (v/v), respectively, and subsequently flash-frozen in liquid nitrogen. Following thawing, samples were centrifuged at 16,000 × g for 10 minutes, and the resulting pellets were resuspended in 500 μL of ice-cold AE buffer (sodium acetate 20 mM, pH 5.2 and 1 mM EDTA). To remove residual DNA, the extracted RNA was treated with RNase-free DNase I (New England Biolabs [NEB], Ipswich, MA, USA) following the manufacturer’s instructions. The quality and integrity of the RNA were evaluated using chip gel electrophoresis with a 2100 Bioanalyzer and RNA Nano chip kit (Agilent Technologies, Santa Clara, CA, USA).

### Bisulfite treatment and m^5^C-Rol-LMAP

Total RNA samples and synthetic RNA oligos were bisulfite-treated using the Zymo EZ RNA Methylation Kit (Zymo Research, Irvine, CA, USA) according to the manufacturer’s instructions. m^5^C-Rol-LAMP was divided in three main steps: hybridization, ligation and PCR amplification. Hybridization reactions were prepared in a total volume of 10 µL containing 3 µL of nuclease-free water, 1 µL of CutSmart Buffer 10x (NEB, Ipswich, MA, USA), 1 µL of 1 µM padlock probe, and 5 µL of bisulfite-treated RNA (100 - 250 ng total). A control without RNA was used as a padlock probe auto ligation control. Reactions were assembled in 0.2 mL PCR strip tubes. Hybridization was performed in a thermal cycler using the following temperature and time profile: 95°C for 1 min, 80°C for 1 min, 70°C for 1 min, 60°C for 1 min, 50°C for 1 min, 40°C for 1 min, and 30°C for 10 min. After the hybridization step, 2 µL of a ligation mixture containing 0.05 U of BST 2.0 DNA polymerase (NEB, Ipswich, MA, USA), 0.5 U of SplintR Ligase (NEB, Ipswich, MA, USA), 12.5 µM ATP, and 5 µM dATP (dGTP or dATP+dGTP) were added to the reaction. The ligation reaction was then incubated in a thermocycler with the following thermal conditions: 37°C for 30 minutes, followed by 65°C for 10 minutes to inactivate the enzymes. Following ligation, samples were diluted by adding 12 µL of nuclease-free water and stored at 4°C. 20 µL RCA-LAMP reaction was set up as follows: 4 µL ligation products, 0.8 µM FIP, 0.8 µM BIP, 100 nM SLP, 1.4 mM dNTP, 6 mM MgSO4, 1× Isothermal Amplification Buffer (20 mM Tris–HCl, 10 mM (NH4)2SO4, 50 mM KCl, 2 mM MgSO4, 0.1% Tween® 20, pH 8.8), 0.16 U/µL Bst 2.0 DNA polymerase (NEB, Ipswich, MA, USA) and 1× LAMP dye (NEB, Ipswich, MA, USA) . Reactions were performed at 65°C, and fluorescence signals were measured every 30 sec (one measurement counted as one cycle) for 1 h. The CFX96TM Real-Time System (Bio-Rad, Hercules, CA, USA) was used for amplification.

### Quantitative analyses

To quantify m⁵C1407 methylation levels in 16S rRNA, a standard curve was generated using a synthetic 41-nucleotide RNA oligonucleotide representing the sequence surrounding position 1407 of *E. coli* 16S rRNA, containing a single 5-methylcytosine (m⁵C) at position 1407. Increasing amounts of this methylated RNA oligo (0.1 ng to 1 ng) were subjected to m⁵C-Rol-LAMP following the previously described method. Ligation was performed in the presence of dATP+dGTP to promote efficient amplification of all correctly circularized padlock probes. Reactions were run in triplicate and the average Ct values were plotted against the log-transformed RNA input amounts to generate a standard curve. Linear regression analysis was performed to assess the correlation between Ct values and input RNA. The resulting standard curve was subsequently used to interpolate methylation percentages in experimental samples by comparing Ct values from dATP-only reactions (detecting unmethylated RNA) with those from dATP+dGTP reactions (detecting total RNA, both methylated and unmethylated).

### Statistical analyses

Differences between pairs of samples were assessed with an unpaired Student’s *t*-test. One-way ANOVA was applied for three group comparisons, followed by Tukey’s post hoc test. Data are presented as mean ± standard deviations of the mean from at least three independent experiments. Data analysis was conducted using GraphPad 10.4.1. A P-value of <0.05 was considered statistically significant. Statistical significance is indicated as follows: *P < 0.05, **P < 0.01, ***P < 0.001, **P < 0.0001; ns, not significant.

## ACKNOWLEDGMENTS

This work was financially supported by the Deutsche Forschungsgemeinschaft (DFG, German Research Foundation): Project numbers 325871075 (SFB 1309) (to KJ) and 464582101 (JU270/21-1 to KJ). S.R.B acknowledges financial support by the Alexander von Humboldt foundation. L.V. acknowledges the support of the Graduate School Life Science Munich (LSM).

